# Self-organizing recruitment of compensatory areas maximizes residual motor performance post-stroke

**DOI:** 10.1101/2024.06.28.601213

**Authors:** Kevin Lee, Victor Barradas, Nicolas Schweighofer

## Abstract

Whereas the orderly recruitment of compensatory motor cortical areas after stroke depends on the size of the motor cortex lesion affecting arm and hand movements, the mechanisms underlying this reorganization are unknown. Here, we hypothesized that the recruitment of compensatory areas results from the motor system’s goal to optimize performance given the anatomical constraints before and after the lesion. This optimization is achieved through two complementary plastic processes: a homeostatic regulation process, which maximizes information transfer in sensory-motor networks, and a reinforcement learning process, which minimizes movement error and effort. To test this hypothesis, we developed a neuro-musculoskeletal model that controls a 7-muscle planar arm via a cortical network that includes a primary motor cortex and a premotor cortex that directly project to spinal motor neurons, and a contra-lesional primary motor cortex that projects to spinal motor neurons via the reticular formation. Synapses in the cortical areas are updated via reinforcement learning and the activity of spinal motor neurons is adjusted through homeostatic regulation. The model replicated neural, muscular, and behavioral outcomes in both non-lesioned and lesioned brains. With increasing lesion sizes, the model demonstrated systematic recruitment of the remaining primary motor cortex, premotor cortex, and contra-lesional cortex. The premotor cortex acted as a reserve area for fine motor control recovery, while the contra-lesional cortex helped avoid paralysis at the cost of poor joint control. Plasticity in spinal motor neurons enabled force generation after large cortical lesions despite weak corticospinal inputs. Compensatory activity in the premotor and contra-lesional motor cortex was more prominent in the early recovery period, gradually decreasing as the network minimized effort. Thus, the orderly recruitment of compensatory areas following strokes of varying sizes results from biologically plausible local plastic processes that maximize performance, whether the brain is intact or lesioned.

## Introduction

The central nervous system displays exquisite reorganization capabilities following lesions caused by stroke. In particular, animal and human studies of recovery of the upper extremity function after stroke have demonstrated a systematic recruitment pattern of compensatory brain areas that depends on the size of the lesions in the cortical motor area (Nudo 2006).

A small lesion affecting the upper limb area of the primary motor cortex has an immediate detrimental effect on the contralateral arm and hand motor performance (Nudo 2007) because the primary motor cortex has extensive projections to spinal motor neurons via the corticospinal tract - about 50% of corticospinal tract projections onto the cervical spine in humans originate in the primary motor cortex (Usuda et al. 2022). Plasticity in the primary motor cortex neighboring the lesion (Nudo and Milliken 1996) has been associated with improved performance, which can return to pre-stroke levels when paired with motor activity (Nudo et al. 1996).

A medium-sized lesion affecting all or large parts of the primary motor cortex controlling the upper extremity leads to a large initial decrease in upper extremity function. However, with recovery, the ipsilesional pre-motor areas reorganize and exhibit increased activity compared to before the stroke; this activity is proportional to the amount of hand representation destroyed in the primary motor cortex (Frost et al. 2003, Fridman et al. 2004, McNeal et al. 2010, Murata et al. 2015, Frost et al. 2022). It has therefore been proposed that the pre-motor areas, via their direct but sparser projections to the motor neuron pools (Maier et al. 2002, Ito et al. 2022, Usuda et al. 2022), can act as a reserve area to control movement formerly handled by the primary motor cortex (Reinkensmeyer et al. 2012, Di Pino et al. 2014).

Finally, a unilateral large-sized lesion affecting most primary and premotor areas leads to initial paralysis in the contralateral arm. However, recruitment of the contra-lesional motor cortex (Cramer et al. 1997, Ward et al. 2003, Bestmann et al. 2010, Rehme et al. 2011, Zaaimi et al. 2012, Murata et al. 2015, Dodd et al. 2017, McPherson et al. 2018, Buetefisch et al. 2023) leads to some degree of recovery of function via ipsilateral projections to the reticular formation, which then projects to the spinal cord via the reticulospinal tract (Illert et al. 1981; see Bradnam et al. 2012 for review). Because reticular projections to the spinal cord are mostly oligosynaptic via propriospinal neurons linking multiple muscles (Peterson et al. 1975, Alstermark et al. 2007), they create muscle synergies in the intact nervous system (Pierrot-Deseilligny 2002). Following a stroke, however, excessive activation of the reticulospinal formation leads to abnormal synergies (Zaaimi et al. 2012, Ellis et al. 2017), which compromise independent joint control during arm movement (Dipietro et al. 2007, McPherson et al. 2018, Li et al. 2019).

Hence, the systematic and orderly sequence of plastic recruitment patterns of compensatory areas following motor cortical lesions of increasing sizes has been well characterized: from the peri-infarct motor cortex to the ipsilesional premotor cortex to the contra-lesional motor areas. What is missing is an understanding of the mechanisms underlying such reorganization. It has recently been proposed that the CNS attempts to increase the descending neural drive to the paretic limb by recruiting “second best” alternative areas (Zaaimi et al. 2012, Choudhury et al. 2019), even if this leads to coarse movements because such movements are preferable to the alternative of paralysis (McPherson et al. 2018).

Here, we generalize and formalize this idea. First, along with current theories of human motor control, e.g., (Todorov and Jordan 2002, Scott 2004, Guigon et al. 2007, Shadmehr et al. 2016, Wang et al. 2016), we assume that the goal of the sensorimotor system is to maximize motor performance, where performance is a function of final error and effort. Crucially, we further assume that this maximization occurs whether the cortical motor system is intact or lesioned. Second, we propose that this performance maximization is achieved conjointly by an unsupervised and a goal-directed reinforcement learning processes. The unsupervised learning process creates the necessary condition for optimizing motor performance because it ensures that neurons fire within their physiological ranges via homeostatic regulation despite large changes in their inputs due to upstream plasticity or lesions. Such homeoplasticity is a feature of sensory and motor neurons both in the intact brain and spinal cord and following central or spinal lesions (Siebner et al. 2004, Murphy and Corbett 2009, Beauparlant et al. 2013, Bains and Schweighofer 2014). As this necessary condition is met, a reinforcement learning process tunes cortical synaptic strengths (or “weights”) to optimize motor performance. Reinforcement learning underlies the learning of motor skills via performance maximization (Gullapalli 1990, Barto 2002, Kambara et al. 2021, Schweighofer 2022). Previous models have shown the feasibility of reinforcement learning in cortical neurons to learn non-trivial motor skills, e.g. (Mazzoni et al. 1991, Reinkensmeyer et al. 2012). Furthermore, reinforcement learning in motor cortical areas is compatible with findings of diffuse dopaminergic inputs to the motor cortical areas (Luft and Schwarz 2009, Hosp et al. 2011) and the role of these inputs in plasticity and motor skill learning (Luft 2009, Molina-Luna et al. 2009).

Based on this theoretical framework, we hypothesize that the self-organizing recruitment of compensatory cortical areas post-stroke attempts to maximize residual upper extremity motor performance given both existing neuro-anatomical constraints in the intact nervous system and new constraints imposed by the lesion.

To test this hypothesis, we built and then systematically lesioned an adaptive neuro-musculoskeletal model that learns to control a simplified planar arm with seven arm muscles. To relate the model to previous experiments in neurotypical and post-stroke human participants (e.g. Dewald et al. 1995, Barradas et al. 2020), we simulated an isometric arm task in which (synthetic) participants generate 2D horizontal forces on a handle attached to the hand.

The neural model contains a primary motor cortex and a premotor cortex that both directly project to spinal motor neurons and a contra-lesional primary motor cortex that projects to spinal motor neurons via the reticular formation. Synapses in the cortical areas are updated via reinforcement learning and the activity of spinal motor neurons is adjusted through homeostatic regulation. We first trained a baseline (non-lesioned) model to reproduce the muscular activity and behavioral performance of neurotypical humans in the task. We then lesioned the model with lesions of different sizes in the primary and premotor cortex. We tracked recovery and neural activity in each region as a function of motor training in early and late recovery periods. Finally, to study the effect of plasticity in each neural area on recovery, we successively blocked the plastic processes in each area (premotor cortex, contralesional motor cortex, and spinal motor neurons).

## Results

### Overview of the model and design of the simulations

We designed and simulated a neuro-musculoskeletal model that comprises a neural network with multiple areas and a simplified linearized musculoskeletal system (Figure 1; see Methods for details). We assumed that the ipsilesional primary motor cortex (M1), ipsilesional premotor cortex (PM), and contralesional motor cortex (CM1) receive desired force information via a (non-modeled) “where” visuomotor pathway. M1 and PM neurons project focally to spinal motor neurons (MN) through the cortico-spinal tract (CST); CM1 projects diffusely by fixed weights to multiple synergic muscles via the reticulospinal tract (RST).

**Figure 1.**
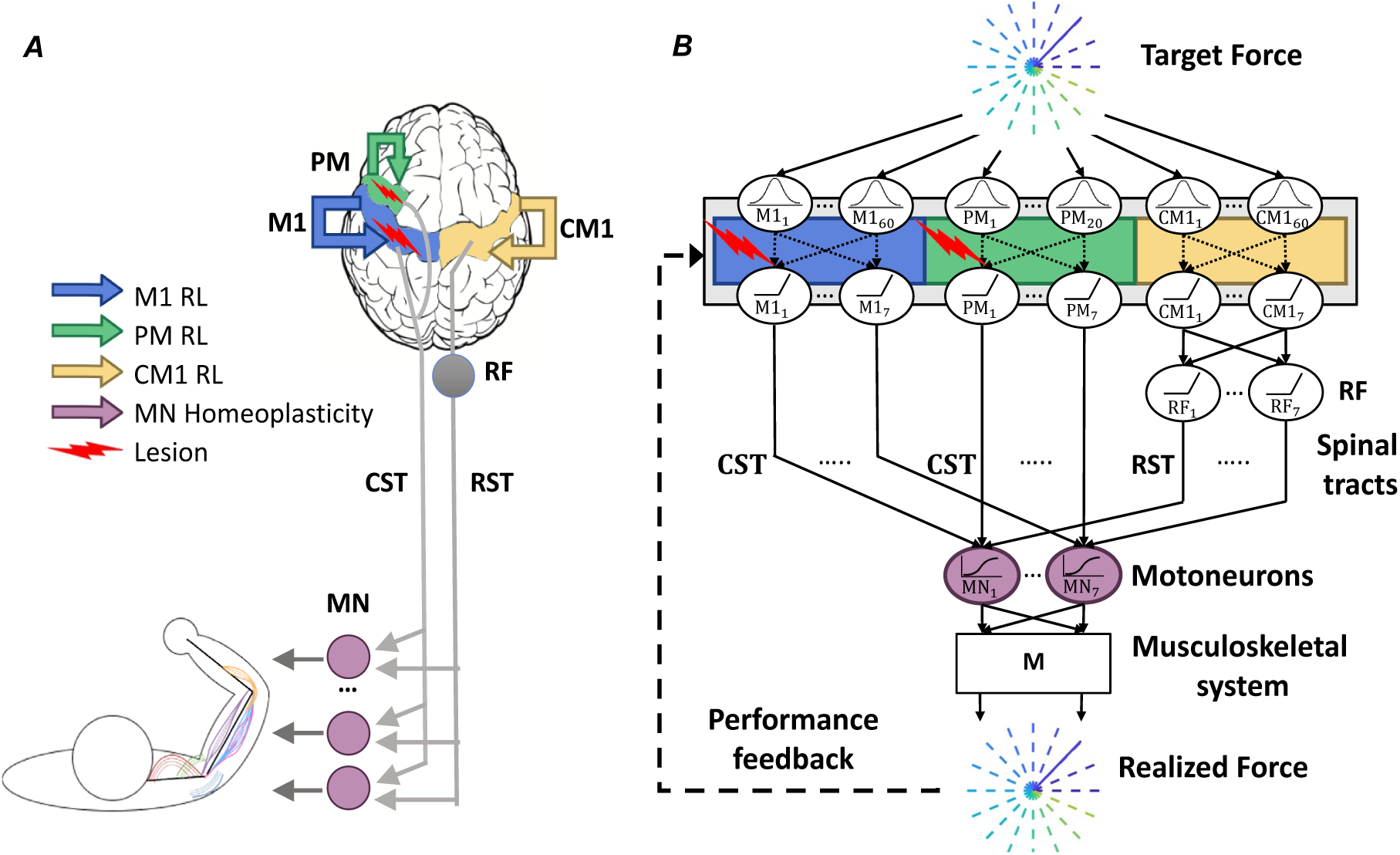
Model Architecture. A. Anatomical diagram showing M1 (blue), PM (green), CM1 (yellow), as sites undergoing reinforcement learning, and MN (purple) as the site undergoing homeoplastic regulation. Lesions (red arrow) of different sizes are simulated in M1 alone or in both M1 and PM. B. Network diagram illustrating the flow of information in the model. Desired force targets are sent to M1, PM, and CM1, which are RBF networks, with fewer RBF neurons in PM than in M1 (see Methods). Neuronal activations from M1 and PM are focally transmitted to MNs via the CST. CM1 projects to the RF, which further projects via synergistic weights to MNs via the RST. Inputs from CST and RST sum in MN sigmoidal neurons, which transmit activity to the data-driven arm musculoskeletal system, implemented with the linear transformation M, which generates the realized force fr. The synaptic weights within PM, M1, and CM1 (shown as dotted lines) are updated via reinforcement learning (RL). The reward depends on force error and effort. MN neurons undergo homeoplasticity that adjusts the sigmoid gain and threshold. M1: primary motor cortex. PM: premotor cortex. CM1: contralesional primary motor cortex. CST: corticospinal tract. RF: reticular formation. RST: reticulospinal tract. MN: motor neurons. M: linear musculoskeletal system. RL: reinforcement learning. RBF: radial basis functions.

We modeled motor cortical neurons (in M1, PM, and CM1) as radial-basis function (RBF) neurons to simplify the neural structure (See Methods for details). We included three times more input neurons in the M1 area than in the PM area to reflect the denser CST projections originating from the primary motor cortex than the premotor cortex. We implemented reinforcement learning in cortical neurons via a local biologically plausible yet theoretically sound stochastic policy gradient rule (Williams 1992) with constant exploration noise. This rule adjusts the weights of cortical RBF neurons to maximize a performance-based reward consisting of a negatively weighted sum of force error and effort (see Methods for details). We then assumed that a homeostatic process regulates the gain and threshold of activation functions in MN neurons to maximize information transfer, as in Triesch (2005) (see Methods and Supplementary Figure S1 for details). Note that we do not model unsupervised learning in cortical neurons for simplicity.

The model simulates an isometric force production task with the arm end-effector, similar to previous experiments with neurotypical and post-stroke subjects, e.g. (Dewald et al. 1995, Berger et al. 2013, Barradas et al. 2020). The goal of the task is to produce a realized force ***f***_r_ at the end-point of the arm, defined here as the wrist, that matches a target force ***f***_t_. The targets ***f***_t_ are contained in a horizontal plane centered at the wrist, and are radially and evenly distributed around the plane’s origin. The musculoskeletal system transforms the output of MN into ***f***_r_ through a linear mapping that represents the action of *n*_musc_ = 7 arm muscles. Such linear mapping has been shown (Zhou and Rymer 2004, Valero-Cuevas et al. 2009, Berger et al. 2013): to provide a good approximation of the actual musculoskeletal system for isometric tasks in a given posture (Figure 2B; see Methods).

**Figure 2.**
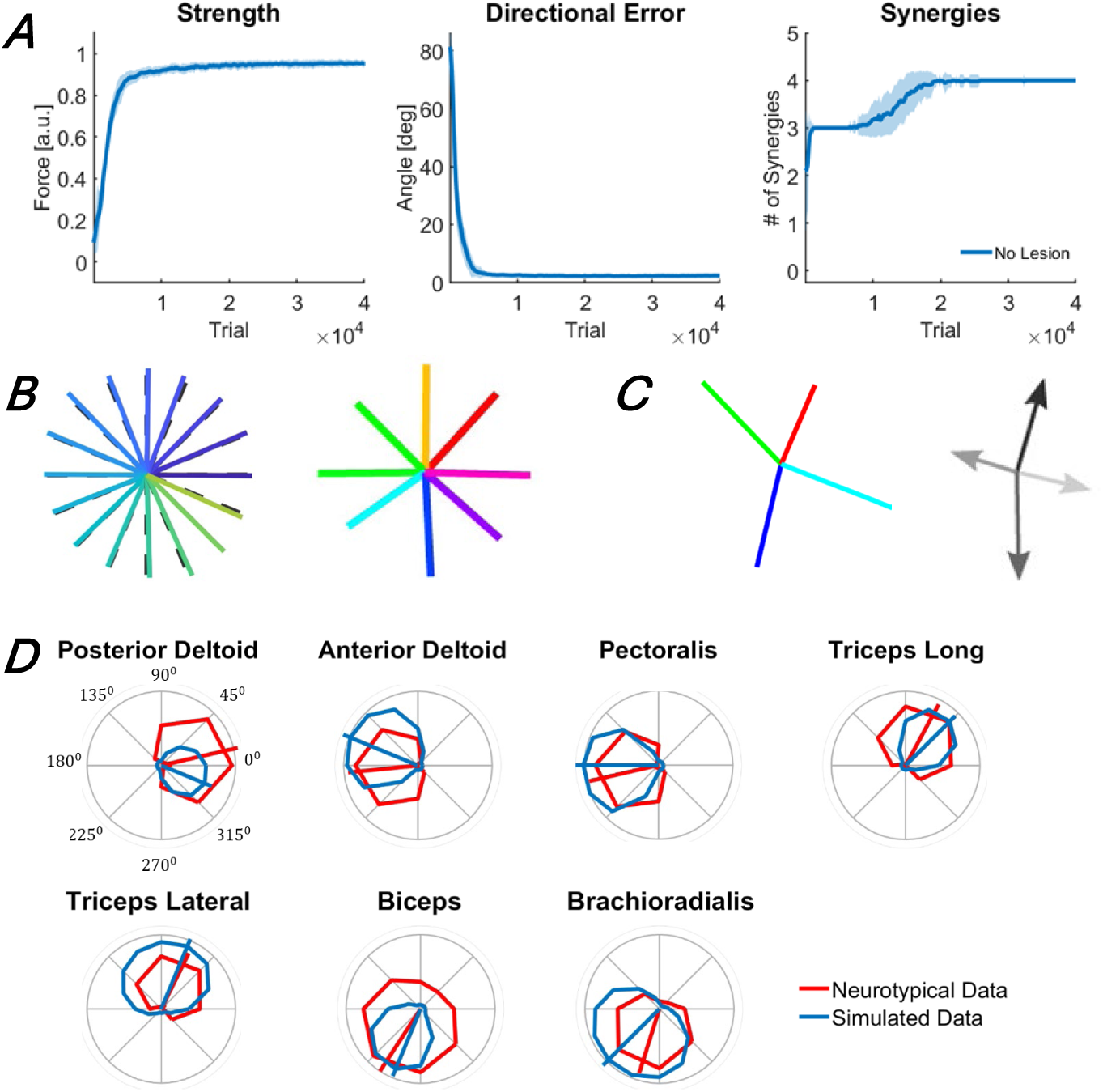
Performance outcomes of the intact baseline model during training (A) and following training (B, C, and D). A. Changes in outcome measures (strength, directional error, and muscle synergies) with training. The traces and shaded areas indicate the mean and standard deviation of the outcome across 50 simulations. B. Target forces (dashed lines) and realized forces (solid lines), produced by the trained model in the isometric arm reaching task to 24 targets (left), and for comparison, forces produced by a representative neurotypical individual (right) in an isometric arm reaching task to 8 targets. C. Forces associated with muscle synergies generated by the model after training (left), and for comparison, muscle synergy forces generated for the representative neurotypical individual (right). D. Muscle tuning curves. Solid blue and red lines indicate the muscle tuning curves of the model after training and those of the representative neurotypical individual, respectively. The segments show the preferred direction of each muscle – all data were collected in the study by Barradas et al. (2020).

We then lesioned the model with lesions of different sizes in M1 and PM, with the lesions destroying a random set of input neurons (by setting their output to 0). We tracked behavioral recovery as a function of motor training via three performance metrics: strength, fine motor control as measured by directional errors, and synergistic muscular activity as measured by the number of synergies, which are extracted via non-negative matrix factorization (NNMF), as used with experimental data (see Methods). To relate the motor deficits and recovery to those of actual patients poststroke, we summarized recovery according to four stages, simplified and adapted to the isometric arm task from the stages proposed by Brunnstrom (Brunnstrom 1966): 1) flaccidity, 2) weak movements and appearance of abnormal synergies, 3) return to normal performance and decrease of abnormal synergies, and 4) complete recovery. We also tracked neural recovery with the sum of neural activity in each area (M1, PM, CM1, and MN). Finally, to test the effect of plasticity in each non-M1 neural area (PM, CM1, and MN), we blocked the plastic processes in each area one at a time. We compared the recovery of each blocked model to the recovery of the full model with all plastic processes.

### Baseline behavior in isometric arm task similar to neurotypical humans

We first trained an intact (i.e., non-lesioned) model to provide performance and neural baselines before we lesion the model and study recovery. We initialized the model with random weights (see Methods) and performed 50 simulations with different sets of initial weights. The model reached asymptotic performance after approximately 3 * 10^4^ training trials (Figure 2A). Because the realized forces ***f***_r_ generated by the baseline model matched the target forces ***f***_t_ almost perfectly, the strength and fine motor control outcomes were close to their maximal values, similar to neurotypical individuals (Barradas et al. 2020) (Figure 2B). Four synergies were necessary to account for 90% of the variance in muscle activations (Figure 2A), and the force vectors associated with these four synergies resemble those of neurotypical individuals in this task (Barradas et al. 2020) (Figure 2C). Finally, the muscle tuning curves obtained from the model were also similar to those of neurotypical individuals for all muscles (Figure 2D), with cosine-like directional tuning and minimization of co-contraction, indicating both realistic muscle directional control for each muscle and adequate effort minimization in the model^1^.

### Recovery and recruitment of brain areas according to stroke lesion size parallel experimental data

Following asymptotic training of the intact model (see above), we made lesions of three different sizes after 4*10^4^ training trials (Figure 3): 50% M1 lesions, 100% M1 lesions, and 100% M1 and PM lesions. We then retrained the models to asymptotic performance with the remaining neural resources.

**Figure 3.**
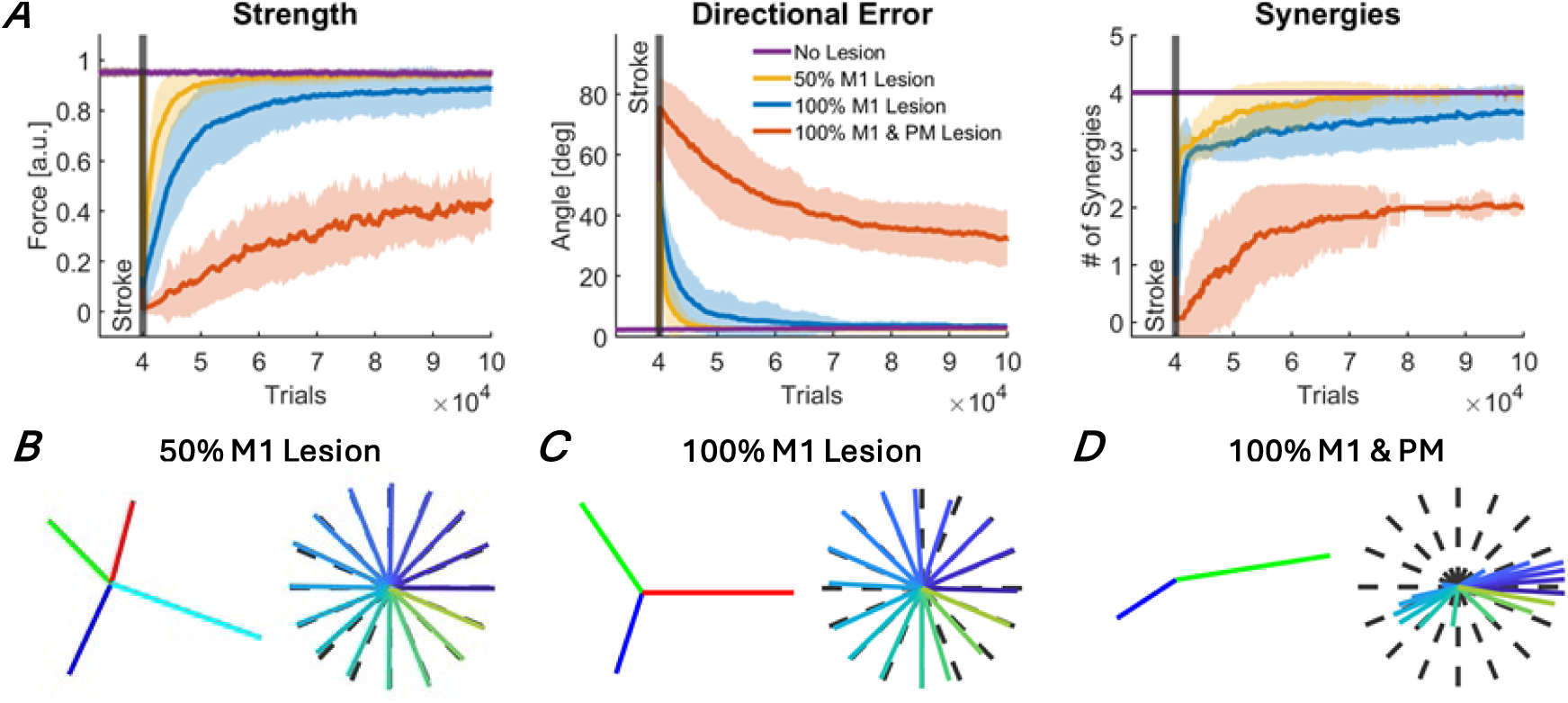
Recovery of motor function after stroke for three different lesion sizes: 50% M1 lesion (yellow), 100% M1 lesion (blue), and 100% M1 and PM lesion (red). A. Recovery of the three outcome measures: strength, directional errors, and muscle synergies. The traces and shaded areas indicate the mean and standard deviation of the outcome across all simulations. The purple trace shows the outcome measures with no lesion. B, C, D. Forces associated with extracted muscle synergies and realized forces at the end of the recovery period. B. 50% M1 lesion. Performance recovered to pre-stroke levels, corresponding to stage 4. C. 100% M1 lesion. Abnormal synergies were still present at the end of recovery, corresponding to stage 3. D. 100% M1 and PM lesion. Weakness, large directional errors, and abnormal synergies were still present at the end of recovery, corresponding to stage 2.

Before stroke, as shown on the left side of the panels of Figure 4 (after asymptotic training of 4*10^4^ trials), the sum of activity in M1 output neurons is large, whereas the sums of activity in PM and CM1 are smaller. Thus, good performance in the baseline model is largely achieved via M1, with both PM and CM1 showing relatively little recruitment.

**Figure 4.**
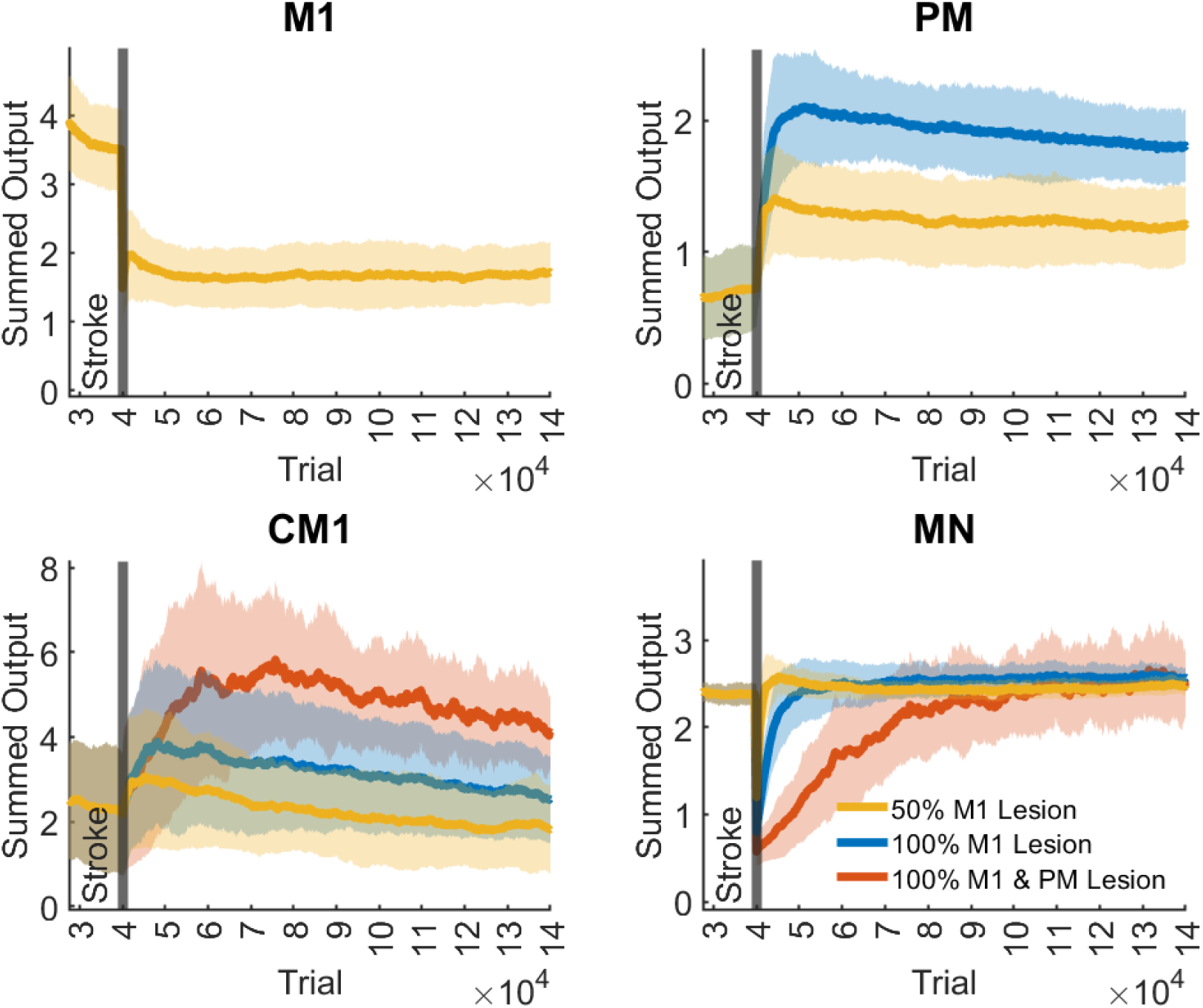
Output activity of M1, PM, CM1, and MN neurons following a 50% M1 lesion, a 100% M1 Lesion, and a 100% M1 & PM Lesion. The output activity is quantified as the sum of all outputs of each region. The traces and shaded areas indicate the mean and standard deviation of the summed output activity across all simulations. Stroke occurred at 4*10 trials, the end of training of the baseline model (Figure 2)

After stroke, all lesions produced a large instantaneous decrease in the performance of the isometric task in all three metrics (strength, directional error, and number of synergies) – see Figure 3. However, the lesion size determined the initial and final performance post-lesion, the rate of recovery, and the number of muscle synergies. For the smallest lesion (50% M1 lesion), the strength, directional error, and synergy number outcomes returned to near prelesion levels after recovery (Figure 3A): the model was able to reach all targets precisely, with as much force as before the lesion, and with three to four synergies, depending on the simulation run. When the model recovered with four synergies, the forces associated with each synergy closely matched those of the original synergies (Figures 2C and 3B). Thus, for the small lesion, the remaining neural resources are sufficient to achieve full recovery, corresponding to stage 4 in our modified Brunnstrom scale.

For a larger lesion (100% M1 lesion, no PM lesion), strength and directional error returned near their pre-lesion levels at the end of training (Figure 3A), with slight deficits in the strength and direction outcomes (Figure 3C). However, muscle synergies remained abnormal at the end of recovery, as shown by the reduced number of synergies compared to prelesion levels, from 4 to ∼3 on average (Figures 3A, right and 3C). Thus, recovery following this lesion is consistent with stage 3 in our modified Brunnstrom scale, i.e., a return to regular movement and abnormal synergies recovery.

For the largest lesion (100% M1 and PM lesion), strength, directional error, and the number of synergies all recovered to only around half of their prelesion levels (Figure 3A). At the end of recovery, only two synergies emerged, which roughly point to the left and the right of the workspace (Figure 3D). As a result, forces could not be generated for upper and lower targets, producing directional errors of around 40 degrees on average for all targets (Figure 3A). These force generation patterns are consistent with observations of movement directions highly biased towards the left and right following large strokes (Reinkensmeyer et al. 2002). Thus, following a complete lesion of the ipsilateral cortex, reinforcement learning in CM1 and homeoplasticity in MN only allowed recovery up to stage 2 in our modified Brunnstrom scale (weakness and emergence of abnormal synergies). In summary, the three lesions of increasing sizes led to a recovery that worsened from stage 4 (50% M1 lesion) to 3 (100% M1 lesion), to 2 (100% M1 and PM lesions).

To study the compensatory roles of the different areas (M1, PM, CM1, and MN) in motor recovery, we quantified the output activity of each area as the sum of all output activations in that area during recovery (Figure 4). After the 50% M1 lesions, M1 showed a nearly two-fold decrease in total activity, as expected. In contrast, PM showed approximately a two-fold increase in activity in the early recovery (after about 2*10^4^ trials post-stroke). CM1 showed an approximate 20% initial increase in activity. However, CM1 activity decayed and returned to near pre-stroke level in the late recovery period (10 * 10^4^ trials post-stroke). MN neurons, after an initial drop, recovered quickly to pre-stroke levels.

After the 100% M1 lesion, both PM and CM1 showed large increases in output activity compared to pre-lesion levels (Figure 4B, C). The increase in PM and CM1 activity in the early recovery period was about three and two times that of baseline, respectively, but decreased in both areas with additional training, most notably in CM1, where activity almost returned to baseline levels. Motor neuron activity showed approximately a two-fold immediate reduction following the lesion but recovered in about 2*10^4^ trials following the lesion. Thus, following a complete lesion of M1, early recovery is driven by large increases in PM and MN activity and to some extent increase in CM1 activity. However, late recovery and final improvements in performance are associated with a relative decrease in activity in PM and CM1 areas, with CM1 activity returning to near baseline levels.

After a complete lesion of the ipsilateral motor cortex (100% M1 and PM lesion), CM1 activity increased to levels between two and three times that of baseline (Figure 4C). The increase was relatively slow compared to changes following smaller lesions, reaching its maximum value after about 3*10^4^ trials post-stroke. CM1 activity then decreased with additional training, reaching about two times that of baseline in late recovery. MN activity dropped about five-fold following the stroke, resulting in practically null strength and, therefore, complete paralysis (see left panel, Figure 3A). However, MN activity fully recovered, albeit at a slow rate in about 5*10^4^ trials, yielding slow recovery in force and directional error following this lesion size (Figure 3A).

The initial large increase in PM and CM1 activity in the early recovery period (at 2 to 3 * 10^4^ trials post-stroke) followed by the decrease in activity in the late recovery period (at 10* 10^4^ trials post-stroke) and the magnitude of these activity levels relative to pre-stroke, mirror those seen in real brains. This is best seen by comparing Figure 5, in which we projected on a brain template the average sum of neural activity during the isometric task after the 50% M1 and 100% M1 lesions in the early and late recovery phases, with PET images following M1 lesion in monkeys (Figure 2 in Murata et al. 2015).

**Figure 5:**
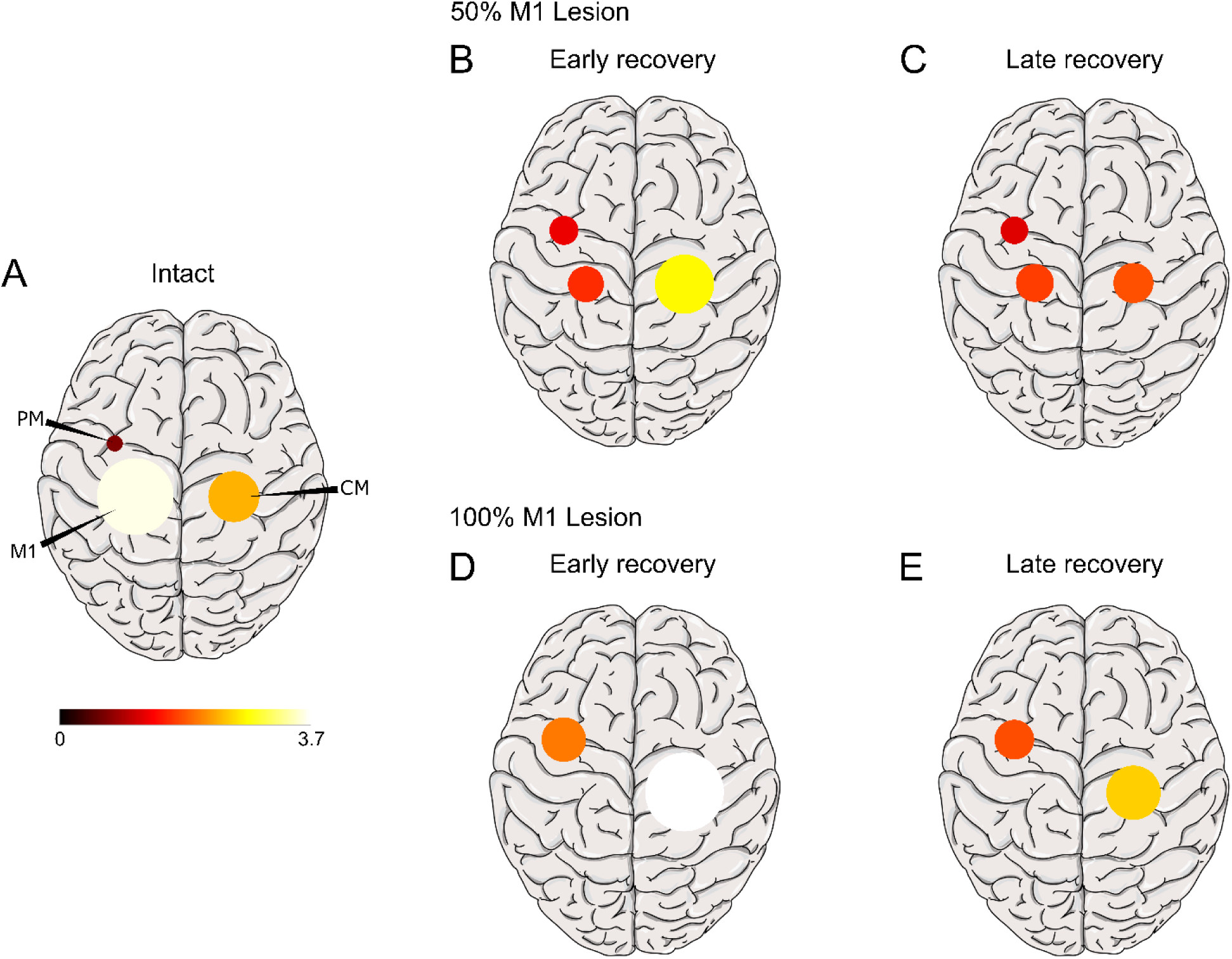
Projection of average sum of neural activity during the isometric task on a brain template in the intact brain (A), and in the early (B, D) and late (C, E) recovery phases following a 50% M1 lesion and a 100% M1 lesion. Early recovery is taken at 10^4^ trials post-stroke and late recovery at 10^5^ trials. Maximum activation is scaled to the maximum recorded. Compare with Figure 2 in Murata et al. 2015.

Why do PM and especially CM1 show an initial stronger activation in the early recovery period and a decrease to lower levels in the late period? As a reminder, the reinforcement learning rule that mediates cortical plasticity tries to maximize a reward that penalizes force error and effort, where high effort is relatively more tolerable than large errors (as expressed by the weight coefficient of effort in the reward). Soon after the stroke, strength is low, generating large errors. PM and CM1 are, therefore, recruited to decrease this error. The recruitment of PM and CM1 also increases effort as MN activity recovers in parallel. In the early recovery period, because the error is large, the effects of high effort on the reward are relatively small, so the contributions of PM and CM1 steadily increase to reduce error. However, as the error starts to be minimized, large efforts become relatively more weighted in the reinforcement learning process. Because the output of CM1 leads to highly synergistic movements associated with large errors (Figure 3D), PM takes over by generating more refined motor commands that produce smaller errors, allowing a reduction of effort in PM and especially CM1 in the late recovery period. This is supported by eliminating the penalties associated with effort in the reward function (effort term = 0 in reward function, equations 8-10), as shown in Supplementary Figure S2. In this case, the activities in PM and CM1 following lesions keep increasing. Thus, when the full reward function was used, as needed to maximize performance and minimize effort, the initial increase in PM and CM1 compensatory areas allows for the quick improvement in reaching performance; this, however, comes at the cost of large effort. Effort is then reduced by further refinements of the synaptic efficacies in these areas, yielding decreasing activities in the late recovery period.

### Plasticity in PM, CM1, and MN differentially impact recovery after a severe stroke in M1: blocking experiments

The above results show that PM and CM1 contribute to recovery following M1 lesions, but their respective contributions to overall recovery and the different motor outcomes are unclear. Therefore, we studied the role of localized plasticity on recovery by blocking plasticity in each area in turn, leaving all other plastic processes active.

When PM plasticity was blocked for the smallest lesion (50% M1 lesion), motor function outcomes returned to near prelesion levels (Figure 6A), reaching stage 4 (full recovery). Thus, for smaller M1 lesions, M1 is sufficient for recovery. Interestingly, in the full model with PM plasticity, there was a significant increase in PM output (see Figure 4). However, the blocking result shows that this “reserve” capacity of PM is not necessary for full recovery after this small lesion. Thus, the reserve effect of PM occurs following a smaller 50% M1 lesion via a cooperative facilitating effect but is not necessary. For the medium lesion (100% M1 lesion), strength, directional errors, and muscle synergy outcomes only reached a fraction of their pre-lesion levels, similar to changes in the full model with 100% M1 and PM lesions (which is unsurprising, as in the full model and the PM-blocked model, CM1 and MN plasticity are the only active processes that can drive recovery). This shows that PM is only truly needed for recovery of fine motor control for large M1 lesions. Finally, we note that performance following the large 100% M1 and PM lesion is worse than the performance after 100% M1 lesion with no PM plasticity; this indicates that undifferentiated PM activity acts as a baseline input to MN neurons that improve performance, notably strength.

**Figure 6.**
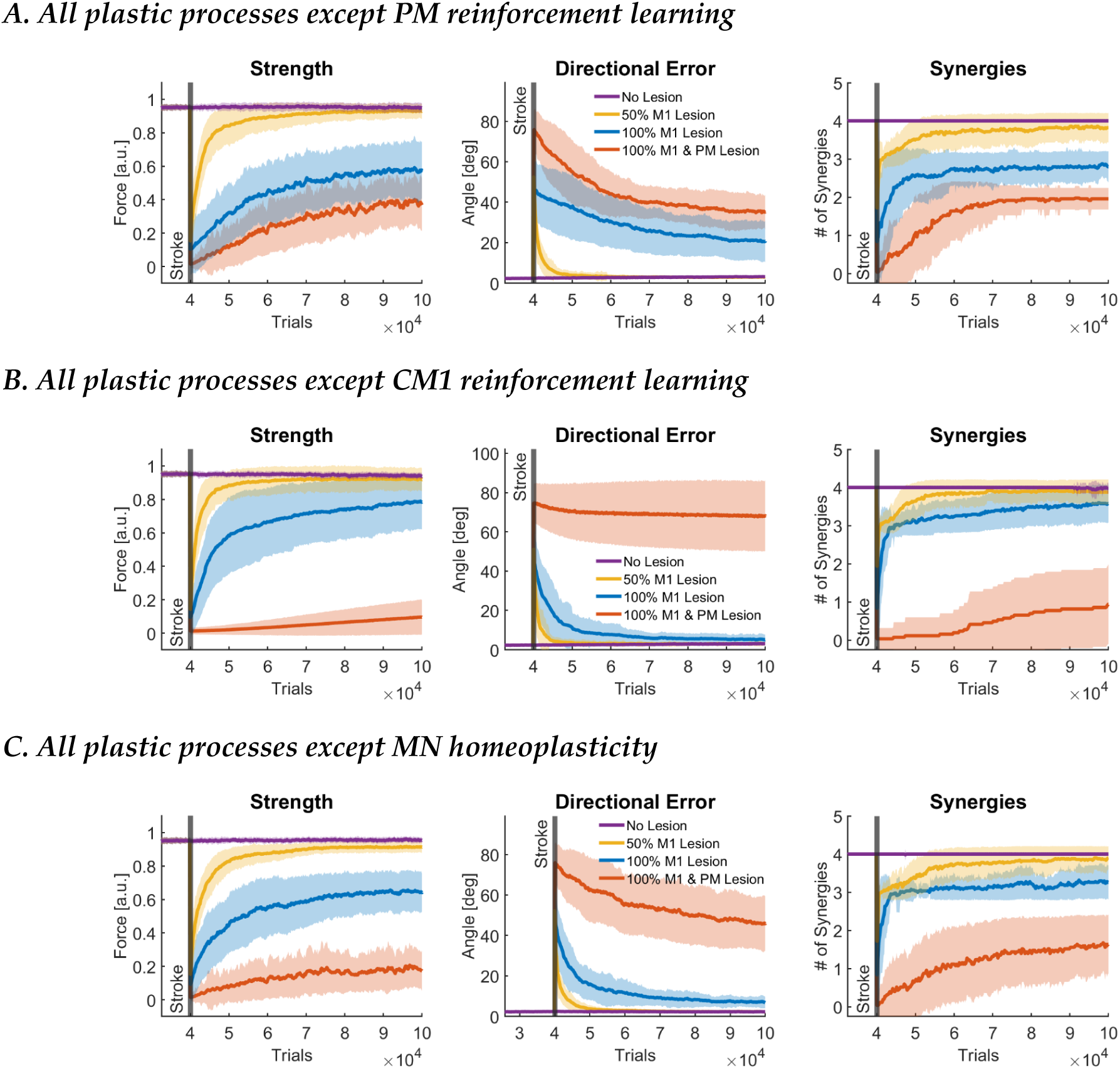
Results of learning blocking experiments. A. Recovery of motor function after stroke with all plastic processes except PM. B. Recovery of motor function after stroke with all plastic processes except CM1. C. Recovery of motor function after stroke with all plastic processes except MN. Data is presented in the same format as in Figure 3A.

When CM1 plasticity is blocked, behavior is similar to the case when all plastic processes are active for both 50% M1 and 100% M1 lesions (Figure 6B, compare with Figure 3). Thus, as in the case of PM above, the increase in activity seen in CM1 after these lesions in the full model (Figure 4) is unnecessary for recovery. Following complete lesion of M1 and PM, there is no recovery when CM1 plasticity is blocked. MN neurons do not fire, and there is almost complete paralysis (stage 1). Thus, CM1 plasticity allows the CNS to overcome paralysis following a large stroke to M1 and PM, but at the price of poor motor control with the emergence of abnormal synergies (Figure 3).

When MN plasticity is blocked, following the smallest lesion (50% M1 lesion), directional errors returned to prelesion levels but with a slight deficit in strength (Figure 6C). For the medium lesion (100% M1 lesion), there is a large deficit in strength at about 60% pre-stroke levels. For the largest lesion (100% M1 and PM lesion), force output is negligible for all directions for the whole recovery period. Thus, recovery remains in stage 1 of our modified Brunnstrom scale (paralysis).

## Discussion and Conclusion

We developed and tested an adaptive neuro-musculoskeletal model that learns to control a simplified arm via four plastic subnetworks involved in arm movement post-stroke. The intact model’s behavior, muscular activity, and overall activation of the cortical areas are comparable to those of neurotypical individuals. We then lesioned the model with different lesion sizes affecting the primary motor cortex and/or the premotor cortex. The model’s neural, muscular, and behavioral results for different lesion sizes account for the behavior and pattern of neural reorganization observed experimentally in the lesioned brain for increasing lesion sizes. Importantly, behavioral recovery and neural reorganization following the different lesion sizes did not follow a preprogrammed master plan. Instead, recovery and the recruitment of compensatory neural activity are self-organizing and emerge from local plastic processes and their interactions with anatomical constraints before and after the lesion.

Specifically, the sequence of re-organization in the cortical areas based on lesion size was due to the ability of each cortical area to generate movements that maximize residual performance and the ability of spinal cord motor neurons to self-regulate despite large changes in their inputs due to the lesion. In the intact brain, the primary motor cortex and spinal cord can together generate optimal behavior, that is, accurate (i.e., low error) and efficient (i.e., low effort) behavior, with other cortical areas only playing a secondary role in motor command generation. In the event of lesions of increasing size, neighboring areas are recruited. If the lesion in the primary motor cortex is too large to generate optimal behavior, the premotor area is recruited first, acting as a first reserve area thanks to its direct cortico-spinal projections - see (Reinkensmeyer et al. 2012). If this recruitment is insufficient to attain good performance, the premotor and contralesional motor areas cooperate to compensate for the loss of primary motor cortex activity. Thus, when the primary motor area cannot “do the job” for large lesions, the premotor area is recruited. For larger lesions that extend to the premotor cortex (100% M1 and PM), compensatory activity is mostly driven by contralesional motor areas, which results in the emergence of two abnormal synergies, causing poor direction control mostly along the directions of the forces associated to these two synergies, as seen in patients with severe stroke (Reinkensmeyer et al. 2002).

Because of the negative correlation between measures of upper limb function and activity in the contra-lesional motor cortex and reticular formation, it has been previously suggested that contra-lesional motor cortex recruitment is maladaptive, e.g. (Boyd et al. 2017, Grefkes and Fink 2020). In contrast, the results of our model, based on the principle that the CNS continuously aims at maximizing performance given the anatomical constraints existing before and after the lesion, are consistent with previous proposals that the CNS uses “second best” available areas (Zaaimi et al. 2012, Choudhury et al. 2019, see also Rehme et al. 2011). Accordingly, premotor cortex recruitment following smaller lesions and contra-lesional motor cortex recruitment following large lesions are adaptive processes that optimize motor performance.

We note that as recovery progresses following 50% and 100% M1 lesions, the output activity of PM and CM1 gradually decreases following an initial large increase – see Figures 4 and 5. As shown by removing the effort term from the reward function (Supplementary Figure S2), this pattern of activity is consistent with a gradual minimization of effort following faster improvement of reaching performance via minimization of errors. Therefore, our model reproduces the compensatory activity of PM and CM1 after an M1 lesion, as observed in actual brains, where a large activity in these areas in early recovery following stroke is gradually reduced in late recovery (Marshall et al. 2000, Ward et al. 2003, Murata et al. 2015, Volz et al. 2017).

Plasticity in cortical areas and recovery in the model crucially depend on a reward signal that encodes both reaching errors and effort. The REINFORCE learning rule reflects the modulation of long-term potentiation, a Hebbian mechanism, by dopamine in the motor cortex. The term *R* − *R̂^t^* in equation 10 corresponds to a reward prediction error that modulates the Hebbian term *ε*_*A*_*ϕ*_*A*_^*T*^, in which the presynaptic input *ϕ*_*A*_ is multiplied by the post-synaptic noise term *ε*_*A*_, which corresponds to the neurons’ output at the current time minus the mean output. A large body of evidence supports this view by showing the important role of rewards carried by dopamine in motor-cortex mediated motor skill learning. Dopaminergic neurons stemming from the ventral tegmental areas are responsive to reward, and in particular reward prediction errors, and provide major inputs to the motor cortex and premotor cortex areas (Luft and Schwarz 2009, Hosp et al. 2011). Neurons in these motor cortical areas are also responsive to rewards (Ramkumar et al. 2016), as dopamine receptors have been shown to be important modulators of long-term potentiation in the motor cortex, with dopaminergic inputs enhancing motor skill learning in animals (Molina-Luna et al. 2009, Hosp et al. 2011, Rioult-Pedotti et al. 2015), and in neurotypical and post-stroke humans (Floel et al. 2008, Floel et al. 2005). Furthermore, a clinical trial showed that the administration of a dopaminergic agonist, L-DOPA, together with physical therapy, improves overall motor functions (Scheidtmann et al. 2001).

Finally, a crucial component of the reward signal in the model is the effort-based term. This term was introduced on theoretical grounds as the objective function in optimal control models of human arm movements includes both error and effort minimization terms. Effort minimization in the model was linked to the decrease in activity in compensatory areas in the late phase of recovery (as shown by simulations without an effort term). In the model, the effect of large errors on reward is larger than the effect of high effort. Thus, the system prioritizes the decrease of performance errors; when this sub-goal is attained, effort then slowly decreases by refinement of the synaptic projections, leading to reduced activity. A prediction from the model is therefore that the slow decreases in motor cortical activity during recovery are related to a decrease in effort.

Homeoplasticity has been proposed to play a significant role after a lesion in the nervous system (Murphy and Corbett 2009, Nahmani and Turrigiano 2014). Homeoplasticity is ubiquitous in the CNS and acts to maintain stable firing rates and patterns. In response to abnormally low or high activity after lesion, homeoplasticity may thus enhance or decrease excitability, respectively, to restore prelesion firing levels (Murphy and Corbett 2009). We previously showed in a model of lesion of the somatosensory cortex that recovery required both homeoplasticity and Hebbian plasticity (Bains and Schweighofer 2014). The absence of homeoplasticity drove maladaptive plasticity and poor reorganization by causing the activity of many cells to remain close to zero. Although we did not include cortical homeoplasticity in the current model for simplicity, a similar phenomenon was observed here in spinal neurons: without homeoplasticity, spinal motor neurons became inactive following large upstream lesions, which led to low strength or even paralysis. Then, because of the deficient network output, the reinforcement system did not receive adequate rewards, precluding performance maximization and recovery. In the full model, the homeostatic regulation process ensured that the spinal motor neurons become responsive again following large lesions to generate appropriate levels of force, allowing reinforcement learning to occur and maximizing recovery. Our study, therefore, suggests that homeoplasticity post-stroke creates the necessary conditions for reorganization and recovery.

### Future directions

Computational models are, by definition, a simplification of the actual neural systems under study. Here, we included only the necessary details to account for behavior and muscular and neural activity in the non-lesioned and lesioned brain in a simple isometric task. We envision five model extensions that can better account for the recruitment of compensatory areas underlying stroke recovery in actual brains. First, the model did not incorporate intra-hemispheric communication, such as inputs from the premotor cortex to the motor cortex, nor bilateral inter-hemispheric connections via the corpus callosum (Wang et al. 2012). These latter connections, in particular, are thought to play an important role in stroke recovery via their overall reciprocal inhibitory effect, which probably results in even lower performance, both directly via further depression of the remaining activity in the ipsilesional areas and indirectly via decreased activity-dependent plasticity. In addition, such a mechanism could enhance contralesional plasticity, further increasing abnormal synergies. Future modeling work is needed to tease out the importance of these connections in reorganization and recovery. Second, whereas the experimental and clinical evidence reviewed above supports the reinforcement learning approach used in the model, our model did not include the basal ganglia or the cerebellum, which are important areas for motor learning and stroke recovery. In particular, the basal ganglia are a major site of dopaminergic-driven reinforcement learning and play a role in habitual skill expression following extensive training (Doya 2000). Additional research is needed to understand the synergistic roles of reward-based plasticity in the motor cortex and the basal ganglia in recovery post-stroke. In addition, the cerebellum is involved in error-based learning. In our model, the (unsigned) error-based part of the reward is similar to the (signed) error signal used by the cerebellum to improve performance (Doya 2000, Kawato et al. 2011). Here again, more research is needed to understand how error and reward differentially and synergistically affect motor skill learning and recovery from stroke. Third, the model does not incorporate a critical “plasticity window” (Murphy and Corbett 2009), during which plasticity is thought to increase. Here, we assumed that training in the model started in this high plasticity state, in line with animal experiments. It is unclear how motor training after this window will affect overall brain re-organization. Fourth, the synthetic patient in our study is always in “training mode.” A previous stroke recovery model better mimics actual patient behavior by incorporating an arm choice module, which allowed the synthetic patient to choose which arm to use when not forced to use the affected arm during therapy (Han et al. 2008, – see also Hidaka et al. 2012). Understanding the effect of training and reduced arm use post-stroke on neural reorganization could be driven by additional modeling work and studies combining wearable sensors and longitudinal imaging. Finally, besides the lesion sizes, the neurons destroyed by the lesion, the initial synaptic weights, and the random exploration due to neuronal noise, our model did not account for patient variability. Thus, recovery and re-organization in the model follow “specific rules” (Grefkes and Fink 2020) dictated by the remaining brain anatomy on the one hand and the goal of the motor system to maximize performance while minimizing effort on the other hand. Future work will need to shed light on the huge variability in recovery, even for similar lesions (Liew et al., 2023).

Whereas recovery post-stroke is typically studied in animals and humans, our neurocomputational modeling approach (Schweighofer 2022) is complementary because it offers four main advantages in studying the mechanisms underlying the recruitment of compensatory areas post-stroke. First, unlike experimental methods, we could simultaneously study and combine mechanisms operating at widely different spatiotemporal scales (synaptic plasticity, neurons, networks, and musculoskeletal system) into a theoretical framework to account for stroke recovery. Second, we could induce different lesion sizes in any brain area instantaneously and at no cost. Third, we could record changes in activity in all areas and in the motor system at once during the whole recovery process, including areas difficult to record with imaging techniques, such as the spinal cord. Finally, we could “block” the plastic processes in different brain areas in turn to study their causal effect on recovery. As a result, our approach allowed us to show mechanistically that the recruitment of compensatory areas serves to maximize residual motor performance post-stroke.

## Methods

### The Neuro-Musculoskeletal model

#### The cortical areas: M1, PM, and CM1

We used Radial Basis Function (RBF) networks to model M1, PM, and CM1. All neural and muscle activation transients are neglected for simplicity, as in previous models (Han et al. 2008, Reinkensmeyer et al. 2012, Barradas et al. 2020). Each area receives a target force input such that

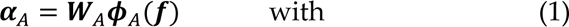

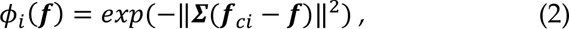

where **α** is the output of the RBF network, ***W*** is a matrix of weights (*n*_musc_ × *n_A_*) that represents the synaptic strength of neurons within each area *A*, ***ϕ***(***f***) is a vector of *n_A_* two-dimensional gaussian RBFs *ϕ_i_*(***f***) evaluated at ***f****, **f***_c*i*_ are the centers of the RBFs, **Σ** is a diagonal matrix with diagonal elements *σ*^2^_RBF_ of equal value, indicating that the RBFs are round, and *A* = {M1, PM, CM1} indicates the represented CNS area. The outputs **α***_A_* of the RBF networks are transformed by a rectified linear activation function to ensure that the inputs to the motoneurons are excitatory:

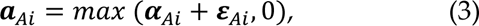

where ***a****_A_* is a vector of excitatory inputs to the spinal motoneurons from cortical area *A*, *i* = 1, …, *n*_musc_ enumerates the output units of area *A*, which for simplicity, we take equal to the number of muscles (see below), and ***ε****_A_* is a vector of exploration noise in the activations of cortical area *A,* such that *ε_Ai_* ∼ *N*(0, *σ*_ε_^2^), and *σ*_ε_^2^ is the variance of the noise. The noise allows plastic processes driven by reinforcement learning, as described in the *Plastic Processes* section.

#### The reticular formation RF

The activations from CM1 project onto the reticular formation and produce outputs ***a***_RF_ that express flexor and extensor synergies:

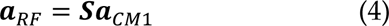

where ***S*** is a matrix of fixed muscle synergies. The synergistic weights are defined to create flexor and extensor synergies and estimated based on muscle pairs that exhibit coactivation. Specifically, the ***S*** matrix is a *n*_musc_ × *n*_musc_ matrix for which the s*_ij_* element is 1 if both muscles *i* and *j* form part of a flexor or extensor synergy, or 0 if not. See below for details about the muscles used in the model. The synergies expressed in RF are responsible for the abnormal co-activation of muscles observed after introducing a lesion in the model (see *Lesion* sub-section).

#### The spinal motor neuron pools MN

The activities within the CST and RST converge at the spinal motor neuron pools. We represent spinal motor neurons as units that sum the CST and RST outputs and transform the sum using a sigmoid function to generate muscle activations ***m***. Each motor neuron pool innervates a single muscle for simplicity:

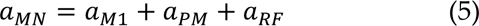

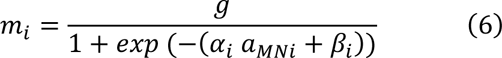

where *i* = 1, …,*n*_musc_ enumerates the output of each motor neuron pool, *g* is a gain that scales the output of the sigmoid, and *α*_*i*_ and *β*_*i*_ are scalar parameters that define the sharpness and threshold of the sigmoid function, respectively. These two parameters are fine-tuned via a homeoplastic process (see *Plastic Processes* section).

#### The musculoskeletal model of isometric force production

Isometric forces produced by the arm in a single posture can be approximated as a linear function of the activations of shoulder and elbow muscles, especially when considering low forces (Zhou and Rymer 2004, Valero-Cuevas et al. 2009, Berger et al. 2013):

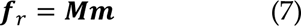

where ***M*** is a matrix (2 × *n*_musc_) that maps muscle activations into planar forces. We modified an experimentally derived linear mapping from a previous study with 10 muscles by selecting a subset of *n*_musc_ = 7 muscles most involved in the horizontal isometric task: posterior deltoid, anterior deltoid, pectoralis major, triceps long head, triceps lateral head, biceps, and brachioradialis. The anterior deltoid, pectoralis major, biceps, and brachioradialis were considered part of the flexor synergy for the definition of the synergy matrix S, whereas the posterior deltoid, triceps long head, and triceps lateral head were considered part of the extensor synergy. Briefly, the mapping was obtained in our previous study by first simultaneously recording EMG and forces and then deriving the mapping via linear regression (Berger et al. 2013, Barradas et al. 2020).

### The Plastic Processes

Our model incorporates four sites of plasticity: M1, PM, CM1, and MN. Plasticity in cortical areas M1, PM, and CM1 is modeled via reinforcement learning that adjusts the synaptic weights ***W***_A_ (equation 1), which allows the acquisition of motor skills based on reward maximization. A loss of motor neurons either in M1, PM, or both leads to decreased input to the motor neurons; such a loss leads to reduced MN outputs and weakness or even paralysis for large lesions. In our model, the MNs can adapt their activation parameters to the input distribution of descending commands through homeoplasticity to maximize information flow through neurons (Bell and Sejnowski 1995, Triesch 2005).

#### Reinforcement Learning in cortical neurons

We assume that the learning processes in the cortical regions M1, PM and, CM1 are driven by reward *R*, which represents a trade-off between force error *e* and effort *u* in the isometric task:

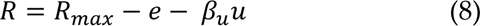

where *R*_max_ is the maximum potential value of the reward, and β_u_ is a factor that weighs the trade-off between error and effort. The error *e* and effort *u* terms are defined as

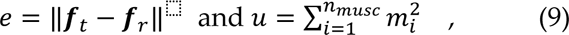

where *e* reflects the model’s accuracy in generating the force to reach the target, and *u* reflects the model’s efficiency in generating the force.

The learning process tunes the synaptic weights ***W****_A_* in the different cortical areas by attempting to minimize the reward prediction error, which expresses differences between the expected reward *R̂* and the actual reward *R* for a given action. We use the simple yet biologically plausible REINFORCE rule, a type of policy gradient rule, to update the synaptic weights ***W****_A_* (Williams 1992):

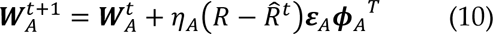

where *η* is a learning rate, **ε** is the vector of exploration noise (equation 3), ***ϕ*** is the vector of RBF representations of the target force (equation 2), the subindex *A* indicates the cortical region of interest, and *R̂^t^* is a reward baseline, given by an exponentially weighted average reward at trial *t*:

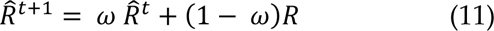

where factor *ω* allows forgetting of previous rewards and learning of new rewards. Finally, we constrain the weights in ***W***_A_ within a range of -*w*_max_ and *w*_max_ and include a momentum term with factor *γ* (not shown in the learning rule) to accelerate learning.

#### Homeoplasticity in MN neurons

For the sigmoidal MNs defined in equation 6, the *α*_*i*_ and *β*_*i*_ parameters that control the sharpness and threshold of the activation function of the spinal neurons are updated according to (Triesch 2005):

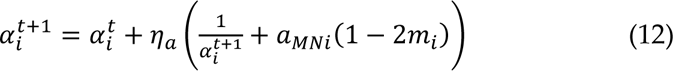

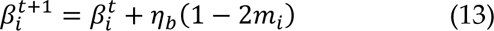

where *η*_*a*_ is a learning rate for *α*_*i*_, *η*_*b*_ is a learning rate for *β*_*i*_, *m_i_* is the muscle activation of muscle *i* (equation 6), and *a*_*MNi*_ is the input to motor neuron *i* (equation 5). These plastic rules thus ensure that the output distribution of individual spinal motor neuron pools after stroke remains equivalent to its pre-stroke distribution despite the decrease of inputs due to the lesion (Figure S1).

### Data analysis

#### The three main outcome measures: force, fine motor control, and synergies

We assessed recovery in the simulations via three outcome measures derived from realized forces ***f***_r_ and muscle activations ***m***: strength, fine motor control, and muscle synergies. Strength measures the ability to generate forces of appropriate amplitudes. In our simulations, we measured strength as the magnitude of fr in each trial. As recovery progresses, the magnitude of ***f***_r_ should approach the magnitude of the target force ***f***_t_, which is set to 1 a.u. (Figure 3A).

Fine motor control is a measurement of the ability to perform accurate movements. In our simulations, we measured fine motor control as (absolute value of) the directional error between fr and ***f***_t_ in each trial. Therefore, as recovery progresses, this measure should decrease.

We used non-negative matrix factorization (nNMF) to identify abnormalities in muscle synergies after the simulated stroke (Cheung et al. 2009). We quantify the abnormalities in the synergies as the number of synergies extracted by nNMF that account for at least 90% of the variance in the muscle activations. In addition, to account for very low forces, if the force associated with a synergy was less than 0.05, then the synergy was set to 0.

Additionally, for the intact network, we compared the simulated and experimental muscle tuning curves by mapping the direction of each of the 16 target forces ***f***_t_ to their corresponding elicited muscle activation (Figure 3E). The experimental data was taken from a previous study (Barradas et al. 2020).

#### Definition of the four stages of recovery based on the three outcome measures

Brunnstrom proposed a sequence of seven stereotypical stages for recovery post-stroke (Brunnstrom 1966) that include flaccidity, emergence of spasticity, increase in spasticity during synergistic voluntary movements, movement patterns outside of synergy and decrease in spasticity, more complex movements with further decrease in spasticity, disappearance of spasticity, and full recovery of coordinated voluntary movements. Because spasticity as commonly defined is movement-related, and thus less relevant to our isometric arm task, we modified and redefined the number of stages of recovery. Stage 1 is characterized by flaccidity. Stage 2 is characterized by weak movements and appearance of clinical movement synergies. Stage 3 is characterized by return to regular movements and decreases of abnormal synergies. Finally, stage 4 is characterized by complete recovery.

### Simulation Procedure

We examined the influence of the four plastic processes described in the *Plastic Processes* section in recovery after cortical stroke lesions of different severity. Each simulation consisted of three phases: 1) baseline phase, 2) lesion phase and 3) recovery phase. For each condition, simulations were repeated 50 times with different random seeds. We present the mean and standard deviation of all outcome measures across all the simulated instances.

#### Baseline phase

In the baseline phase, the model learned to produce behavior that is similar to the behavior of healthy subjects in an isometric arm reaching task (Figure 3). The baseline state of the model was obtained by using reinforcement learning in M1, PM and CM1, and homeoplasticity in the spinal motor neuron pools, as described in the *Plastic Processes* section. The connections and parameters in each component of the model were randomly initialized (see parameter table). A trial in the isometric task consisted of presentation of target ***f***_t_, generation of realized force ***f***_r_, and a learning iteration for all plastic areas. In each trial, the target ***f***_t_ was randomly selected from a set of 16 radially and uniformly distributed targets, such that every 16 trials all targets were selected once. The baseline phase consisted of 40 * 10^3^ trials, although performance in the task stabilized after approximately 30 * 10^3^ trials (Figure 3).

#### Lesion phase

After the performance of the model stabilized at the baseline state, a simulated cortical lesion was introduced. We simulated cortical lesions of different severity levels. The lesions were implemented by nullifying the activations *ϕ_i_*(***f***) of affected neurons (RBF units) in the lesioned area, which is equivalent to removing the affected neurons. We defined the severity of the lesions as the percentage of affected neurons in the lesioned area. We simulated three different lesions: 50% M1, 100% M1, and 100% M1 and PM. In all runs for the 50% M1 lesion, we randomly selected the neurons to be lesioned.

#### Recovery phase

After the lesion, performance in the isometric task is degraded, and training is continued to attempt to restore pre-lesion levels of performance. Trials in the recovery phase follow the same procedure as the baseline phase: presentation of target ***f***_t_, generation of realized force ***f***_r_, and a learning iteration. However, to examine the effect of each plastic process in the model, we delimit the areas of the CST in which learning takes place. We defined 4 post-lesion plasticity conditions: 1) full plasticity (plasticity in M1, PM, CM1 and MN), 2) plasticity in all areas but PM, 3) plasticity in all areas but CM1, and 4) plasticity in all areas but MN. Selectively removing plasticity in each area allowed us to determine the role of each plastic process in promoting recovery. In total, we simulated 12 lesions × plasticity conditions (all combinations of lesion severity and post-lesion plasticity conditions).

#### Parameters

The RBF centers ***f***_c*i*_ are evenly distributed around a circle with an amplitude of 1 (arbitrary units). To represent the denser CST originating from primary than from premotor areas (Usuda et al. 2022), the size of the neuron populations of PM *n*_PM_ is smaller than that of M1 *n*_M1_ and CM1 *n*_CM1_. Furthermore, we initialized ***W***_PM_ and ***W***_CM1_ at 10% of the connection strength of ***W***_M1_. The weights ***W****_A_* were randomly initialized following a lognormal distribution to simulate the heavy tail distribution observed in the brain (Buzsaki and Mizuseki 2014).

| Parameter | Meaning | Value |
| --- | --- | --- |
| $n_{M1}$ | Number of neurons in primary motor cortex | 60 |
| $n_{CM1}$ | Number of neurons in contralesional motor cortex | 60 |
| $n_{PM}$ | Number of neurons in premotor cortex | 20 |
| $n_{musc}$ | Number of motor neurons | 7 |
| $\sigma^2_{RBF}$ | Width of M1, PM, and CM1 RBF Neurons | 0.5 |
| $\sigma_{\epsilon}^2$ | Variance of normal distribution for exploratory noise | 0.03 |
| $w_{mean}$ | Mean of lognormal distribution of initial weights | 0.05 |
| $w_{var}$ | Variance of lognormal distribution of initial weights | 0.0025 |
| $a_{ini}$ | Initial MN sigmoid sharpness | 1 |
| $b_{ini}$ | Initial MN sigmoid threshold | 3 |
| $g$ | MN sigmoid gain | 2 |
| $\eta_a$ | Learning rate for sigmoid slope | 0.00004 |
| $\eta_b$ | Learning rate for sigmoid threshold | 0.00001 |
| $R_{max}$ | Maximum reward | 1 |
| $\beta_u$ | Tradeoff between error and effort | 0.07 |
| $\omega$ | Reward average decay parameter | 0.999 |
| $\eta$ | Reinforcement learning rate in M1, PM and CM1 | 0.001 |
| $\gamma$ | Momentum factor | 0.99 |
| $w_{max}$ | Max weight in $W_A$ | 0.2 |

## Acknowledgments

This work was funded by grant NIH R56 NS126748 to NS.

## Footnotes

1 Note that in the model, the cortical input units also show directional tuning and, for each area, population vector properties pointing in the direction of the target force. These properties emerge from the model structure of the radial basis function neurons with Gaussian tuning, as described in Methods.

