## Supplementary figures for "Self-organizing recruitment of compensatory areas maximizes residual motor performance post-stroke"

### 1. How homeoplastic regulation maximizes information transfer

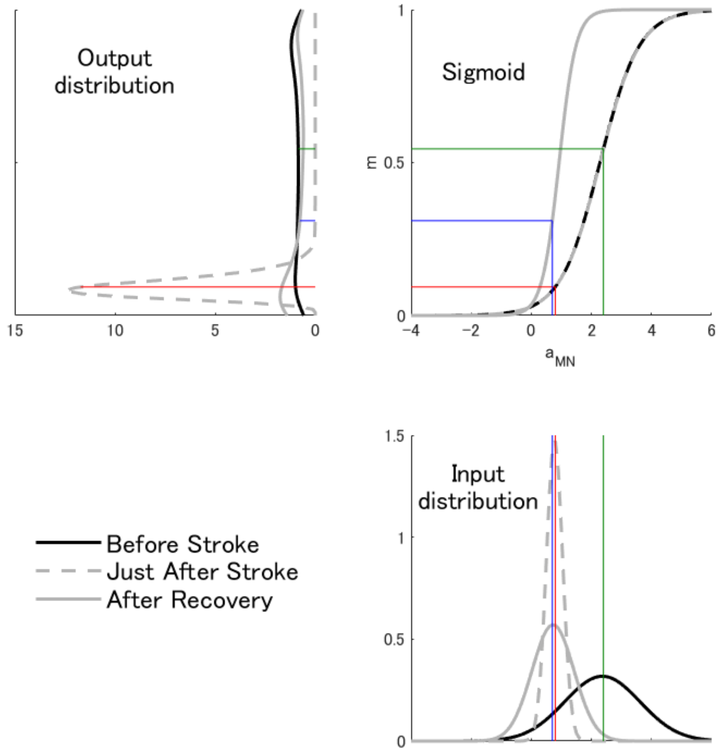


Figure S1. Homeoplasticity in a single spinal motor neuron maximizes information transfer before stroke and after recovery. The input distribution of motor neuron inputs (lower panel) is transformed by the sigmoid (upper right panel) to yield the distribution of motor neuron outputs(upper left panel). Before stroke (black trace), the sigmoidal activation function of the motor neuron pool transforms the distribution of inputs $a_{MN}$ into a flat distribution of muscle activations $m$. However, at the onset of stroke (dotted gray trace), the input distribution shifts to smaller values and becomes narrower due to the loss of input from cortical areas. The unadapted activation function thus produces an output distribution that is highly biased around low values of muscle activation $m$ yielding low strength or even paralysis. During recovery (solid gray trace), homeoplasticity tunes the activation function to restore the output distribution to its pre-stroke shape. The colored lines traversing the input distribution, activation function, and output distribution plots show the mapping of the mean of the input distributions to their corresponding output values. The data in this figure was generated by simulating a 50% M1 lesion and activating only M1 and MN plasticity.

### 2. Activity in compensatory areas without effort minimization

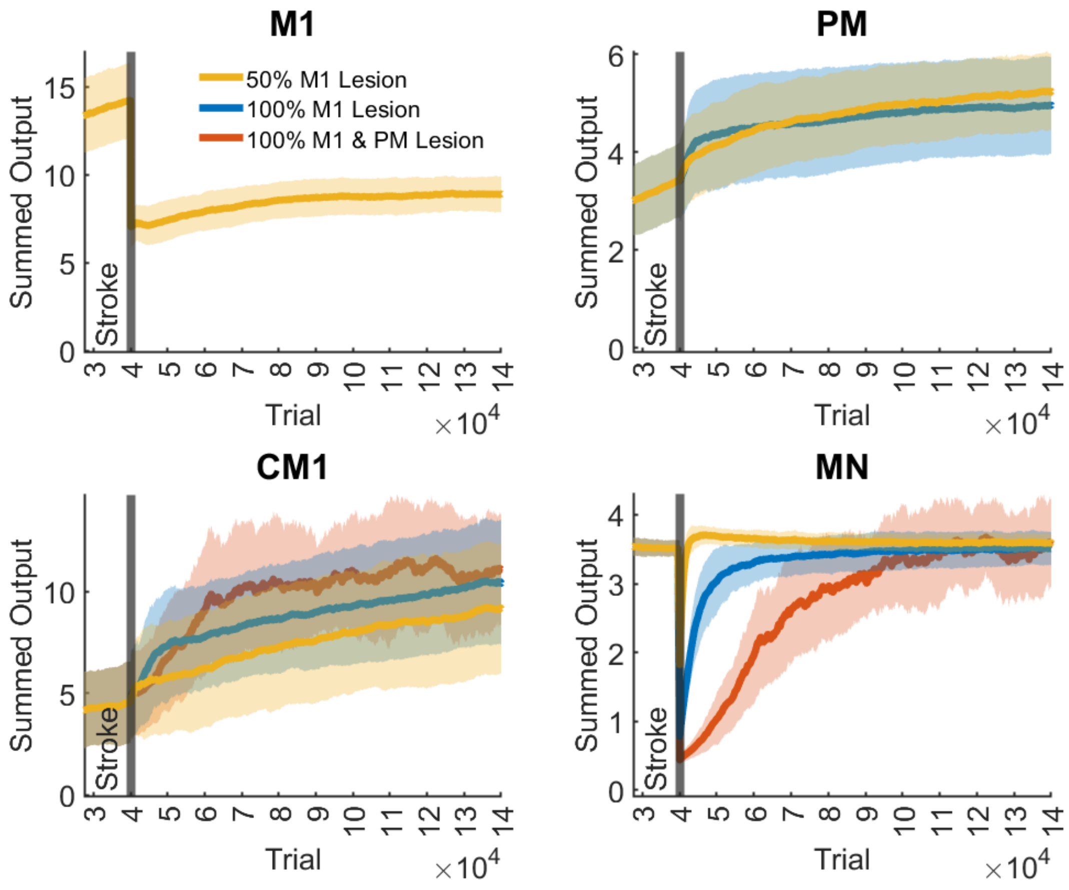


Figure S2: Output activity of M1, PM, CM1, and MN neurons following a 100% M1 Lesion and a 100% M1 & PM Lesion, as in the main text Figure 4, with no effort term in the reward function of equation 8. Note how the activity in PM and CM1 following lesions keeps increasing, whereas in the full model, the activity decreases in late recovery following an initial increase in early recovery. The output activity is quantified as the sum of all outputs of each region. The traces and shaded areas indicate the mean and standard deviation of the summed output activity across all simulations.
